# Enhancer sharing promotes neighborhoods of transcriptional regulation across eukaryotes

**DOI:** 10.1101/061218

**Authors:** Porfirio Quintero-Cadena, Paul W. Sternberg

**Affiliations:** Division of Biology and Biological Engineering, California Institute of Technology; Howard Hughes Medical Institute, Pasadena, California, USA

## Abstract

Enhancers physically interact with transcriptional promoters, looping over distances that can span multiple regulatory elements. Given that enhancer-promoter (EP) interactions generally occur via common protein complexes, it is unclear whether EP pairing is predominantly deterministic or proximity guided. Here we present cross-organismic evidence suggesting that most EP pairs are compatible, largely determined by physical proximity rather than specific interactions. By re-analyzing transcriptome datasets, we find that the transcription of gene neighbors is correlated over distances that scale with genome size. We experimentally show that non-specific EP interactions can explain such correlation, and that EP distance acts as a scaling factor for the transcriptional influence of an enhancer. We propose that enhancer sharing is commonplace among eukaryotes, and that EP distance is an important layer of information in gene regulation.

## Introduction

Enhancers mediate the transcriptional regulation of gene expression, enabling isogenic cells to exhibit remarkable phenotypic diversity (Davidson and Peter, 2015). In complex with transcription factors, they interact with promoters via chromatin looping (Marsman and Horsfield, 2012), finely regulating transcription in time and space. A prevailing view is that most enhancers have a mechanism to selectively loop to a target promoter (van Arensbergen et al., 2014). Examples in this category usually require specific transcription factor binding at both enhancer and promoter sites (Davidson and Peter, 2015), which could explain why some enhancers seem to influence different promoters in varying degrees (Gehrig et al., 2009). On the other hand, EP looping is generally mediated by common protein complexes (Kagey et al., 2010; Malik and Roeder, 2010), conflicting with the specific molecular interactions required by such a model at a larger scale. Examples of non-specific EP pairing seem to be common (Butler and Kadonaga, 2001), yet this model could result in transcriptional crosstalk, which appears inconsistent with our current paradigm of gene regulation. The predominant EP pairing scheme-specific or non-specific-and its determinants are thus unclear. Here we ask to what extent are pontential EP pairs compatible through a metaanalysis of the genome-wide transcription of gene neighbors in five species. We propose that enhancer sharing occurs widely across eukaryotes, test key aspects of this hypothesis in *C. elegans*, and analyze its implications in other genomic phenomena.

## Results and Discussion

### Gene neighbors are transcriptionally correlated genome-wide

To investigate the transcriptional relationship between gene neighbors, we paired every protein-coding gene of five organisms (*Saccharomyces cerevisiae, Caenorhabditis elegans, Drosophila melanogaster, Mus musculus* and *Homo sapiens*) with its 100 nearest neighbors within the same chromosome, which yielded lists of around 600,000 (*S. cerevisiae*) and 2 million (each of the rest) gene pairs. We then computed the Spearman correlation coefficient between paired genes across multiple RNA-seq datasets (Attrill et al., 2016; Ellahi et al., 2015; Gerstein et al., 2010; The ENCODE Project Consortium, 2012) and the intergenic distance between the the start of the 5’ untranslated region of the first gene to the start of the second gene in each pair. We observed that neighboring genes tend to be correlated in transcript abundance genome-wide in all analyzed organisms, and that this correlation decays exponentially with increasing intergenic distance (Figure 1a). We thus fitted the data to an exponential decay function to compute the mean distance at which a pair of genes remain correlated (*d_exp_*). Consistent with the persistence of the correlation pattern across organisms, *d_exp_* scaled with genome size, to 1 kilobase in *S. cerevisiae*, ~10 kb in *C. elegans* and *D. melanogaster*, and ~350 kb in *M. musculus* and in *H. sapiens* (Supplementary Figure 1). This trend remained largely the same even after removing duplicated genes pairs (Supplementary Figure 2). Most genes had at least one neighbor closer than *d_exp_* in all species (Figure 1b), and the representation of gene ontology annotations remained unbiased in correlated gene pairs (Supplementary Figure 3), indicating that the average gene is correlated in expression with its nearest neighbors beyond any particular gene class. In addition, sampled intergenic distances go well beyond *d_exp_* (Figure 1c), indicating that 100 gene neighbors is a sufficient number to study this effect.

**Figure 1:**
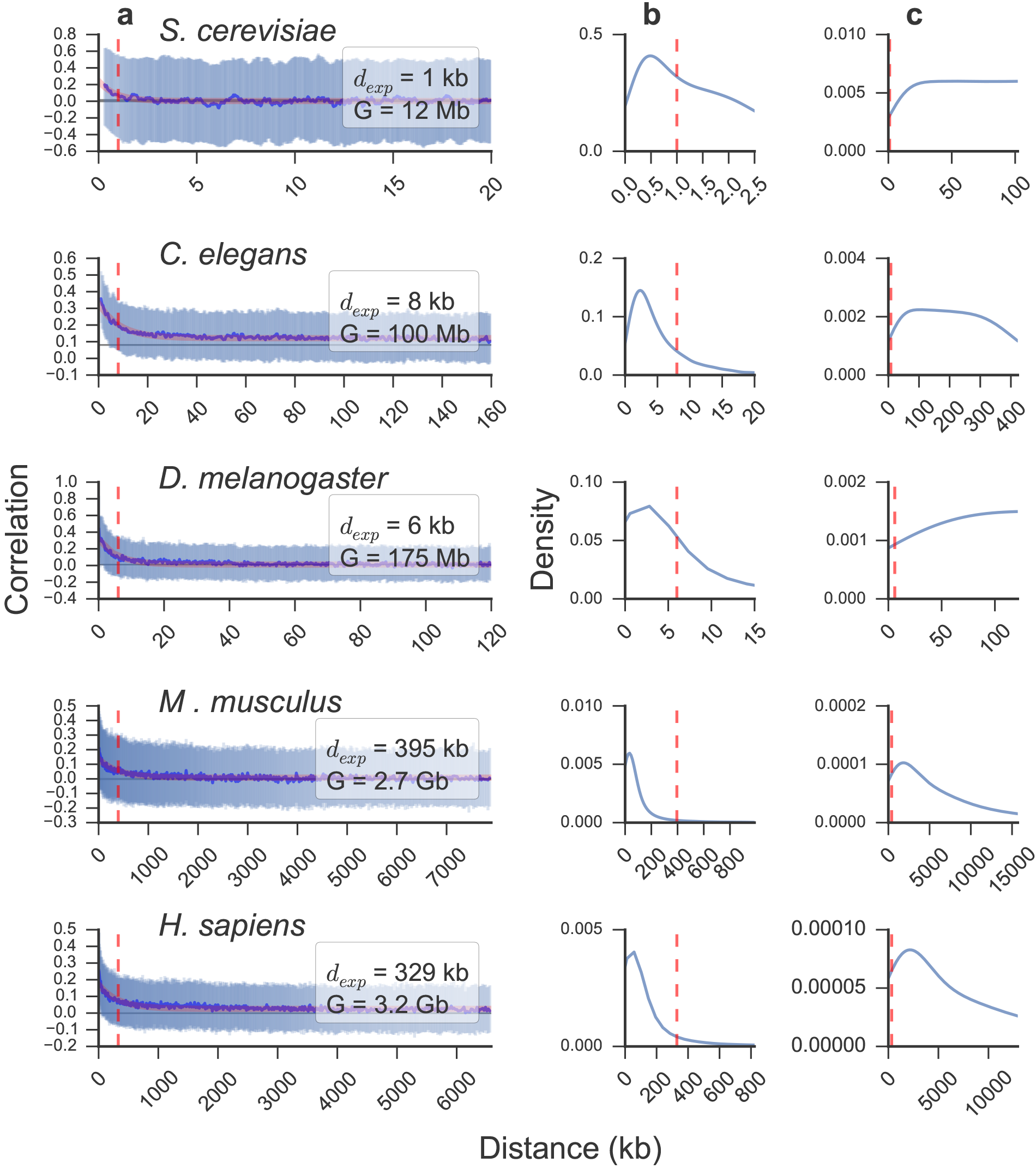
Neighboring genes are transcriptionally correlated genome-wide across eukaryotes. a) Sliding median of correlations between paired neighbors (blue line) and interquartile range (pale blue) with increasing intergenic distance. Median ± 95% confidence interval of randomly paired genes is shown as a horizontal gray line. Fit to an exponential decay function (red line) was used to compute the mean distance at which gene neighbors remain correlated (*d_exp_*, vertical red dashed line). The genome size (G) is displayed for each organism. Distribution of intergenic distances between each gene and its nearest neighbor (b) and all paired genes (c). The organism analyzed in each case is indicated for each group of three subplots.

To examine the correlation of gene expression in the spatial domain, we analyzed RNA *in situ* hybridization data for 6053 genes in *D. melanogaster* (Hammonds et al., 2013; Tomancak et al., 2002, 2007). We computed the percentage overlap in tissue expression by dividing the number of common tissues over the total number of unique tissues in which genes of any given pair are expressed (Supplementary Figure 4a). This analysis revealed that close neighbors have a tendency to be expressed in the same tissues, and that this overlap also decays exponentially with intergenic distance (Supplementary Figure 4b). However, the correlation extends to a longer mean distance (*d_exp_* = 22 compared to 6 kb), suggesting that RNA-seq analysis, which included mostly whole-organism transcriptome averages, resulted in a conservative estimate. Gene neighbors thus have a spatio-temporal correlation in expression that is highly dependent upon the spacing between them. Our meta-analysis extends the findings of previous reports (Michalak, 2008) genome-wide and across organisms. In particular, pairing every gene with 100 proximal genes provides a complete set of distance-dependent correlations between gene pairs.

### Enhancer sharing explains the transcriptional correlation of gene neighbors

The pervasive nature of proximal gene co-expression suggested a common underlying mechanism. We hypothesized enhancer sharing among nearby genes to be a plausible explanation, as it is in agreement with several observations: i) enhancers regulate transcription by making contact with promoters via chromatin looping (He et al., 2014), whose incidence also decays exponentially as the distance between contacting sites increases (Rao et al., 2014; Ringrose et al., 1999), with the same pattern as observed here (Supplementary Figure 5) ii) the average distance between studied EP interactions scales with genome size in ranges consistent with *d_exp_* for each analyzed organism (Araya et al., 2014; He et al., 2014) iii) broad enhancer compatibility and sharing is consistent with the idea that common protein complexes such as the mediator are widely utilized bridges in EP looping (Kagey et al., 2010; Malik and Roeder, 2010) and iv) a high frequency of chromatin interactions are observed within topologically associated domains identified through high-resolution Chromosome Conformation Capture (Hi-C) (Rao et al., 2014). Consistent with this view, it seems that enhancers often do not show promoter specificity (Butler and Kadonaga, 2001). This line of reasoning suggests a model where, as opposed to only having a specific target gene (Figure 2a), enhancers have a range of action in which they can influence any gene (Figure 2b), at least to an extent that causes the correlation of nearby genes, likely within a topological domain. Transcriptome analysis could thus provide indirect evidence of genome and condition-wide EP looping that is difficult to access through Hi-C (Rao et al., 2014) due to the low signal-to-noise ratio of short-range interactions.

**Figure 2:**
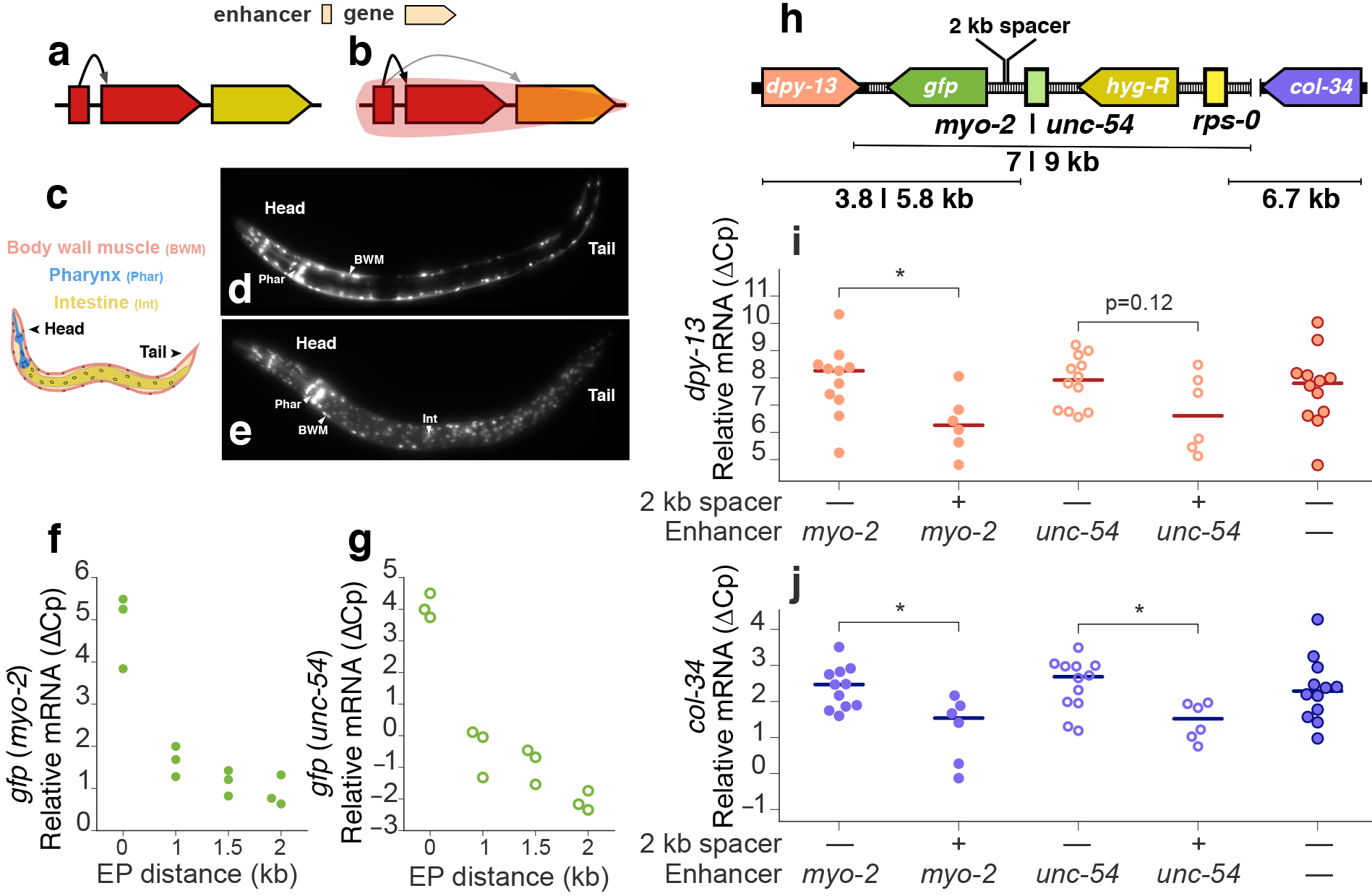
Enhancer sharing explains the transcriptional correlation of gene neighbors. Two possible models for EP relationship: a) Enhancers have specific target genes and b) enhancers have a range of action in which they influence genes by physical proximity. Tissue specific enhancers (c) are generally compatible. Pharynx and body wall muscle (d) and pharynx, body wall muscle and intestine (e) enhancers driving nuclear *gfp* expression. mRNA levels of *gfp* with increasing EP distance for lines with *myo-2* (filled circles, f) and *unc-54* (hollow circles, g) enhancers. h) Genomic context of the integration site. The inserted construct is shown over a dashed black line and includes a hygromycin resistance gene (*hyg-R*) regulated by a ribosomal enhancer (*rps-0*) and promoter in addition to the reporter (*gfp*) regulated by either the *myo-2* or *unc-54* enhancers; the native genes *dpy-13* and *col-34* flank the insertion site. Relative mRNA levels of *dpy-13* (i) and *col-34* (j) in wild-type and lines with and without the 2 kb spacer (*two tailed P-val<0.05, Mann-Whitney U test). The difference in crossing point-PCR-cycle (ΔCp) with the reference gene *pmp-3* and the corresponding median for each group of biological replicates is shown for every qPCR experiment.

Because of its compact genome, rapid development and availability of tissue specific enhancers (Corsi et al., 2015), we decided to use *C. elegans* to test this hypothesis. We first postulated that unrelated enhancers should generally be compatible, showing qualitative additivity when placed upstream of a single promoter. We thus paired the well characterized *myo-2* pharyngeal enhancer with a body wall muscle (BWM) and a BWM plus intestine enhancer, placed them upstream a minimal promoter and a *gfp* reporter, and examined expression in transgenic animals. In both cases, we observed fluorescence in the corresponding tissues (Figure 2c, d, e). This observation is consistent with typical enhancer studies in artificial constructs (Dupuy et al., 2004) and agrees with some EP compatibility studies (Butler and Kadonaga, 2001).

Given that both chromatin looping and expression correlation decay exponentially, we reasoned that transcription of a given gene should also decay exponentially with increasing EP distance if the observed correlation is to be explained by enhancer sharing. To test this hypothesis, we first built a series of genetic constructs with increasing neutral EP distances (0, 1, 1.5 and 2 kb) for two different enhancers, *myo-2* and *unc-54*. We then integrated each construct in single copy into the genome of *C. elegans* and used quantitative PCR to i) measure the influence of EP distance on the reporter gene in native chromatin and ii) analyze the impact of the perturbation on the two genes that natively flank the site of transgene insertion (*dpy-13* and *col-34*, Figure 2h), which we reasoned should be affected in two counteracting ways. First, the ectopic enhancers should promote their expression. Second, the increased EP distance caused by the addition of spacers should reduce their expression by scaling down the influence of both ectopic and native enhancers (the latter of unknown identity and location) to the opposite side of the spacer.

We found that transcriptional levels of the reporter gene indeed fall rapidly with increasing EP distance with both enhancers (Figure 2f, g); this occurred at a rate that seems congruent or faster than the genome-wide correlation decay, likely because of the dramatic separation of every regulatory element at once, as opposed to gradual separation from individual enhancers in a native environment. Transcription was still well detected even when the enhancers were placed 2 kb away, supporting the hypothesis that EP distance is a scaling factor on the enhancer’s influence. Expression of *dpy-13* and *col-34* was reduced with the introduction of the 2 kb spacer when compared to transgenic lines without it (Figure 2i, j). On the other hand, spacer-free lines were comparable to wild-type, suggesting the incorporation of ectopic enhancers compensated for the EP distance increase caused by the addition of the genetic construct itself. Hence, these observations fit the corollaries of our model even amid the complexity of a native regulatory environment.

### Enhancer-promoter distance insulates neighboring genes

We next wished to determine the extent to which enhancer sharing explains other genomic phenomena. Previous reports have suggested that divergent, parallel and convergent gene pairs tend to have distinct correlation profiles (e.g. (Chen and Stein, 2006)). To explore this hypothesis, we first compared the distribution of intergenic distances of gene pairs oriented in parallel, divergent and convergent orientations (Figure 3a, Supplementary Figure 6). As expected, divergent gene pairs tend to be closest, followed by parallel and finally convergent genes. We then confirmed that each group appears to have different distributions of correlations (Figure 3b, Supplementary Figure 6). To consider the influence of EP distance, we sampled gene pairs from each orientation controlling for intergenic size. This resulted in distributions of correlations that exactly overlap (Figure 3c, Supplementary Figure 6), an observation that is supported by previous reports (Ghanbarian and Hurst, 2015). We thus conclude that the apparent influence of gene orientation in the transcriptional relationship of neighboring gene pairs is a consequence of enhancer sharing and EP distance.

**Figure 3:**
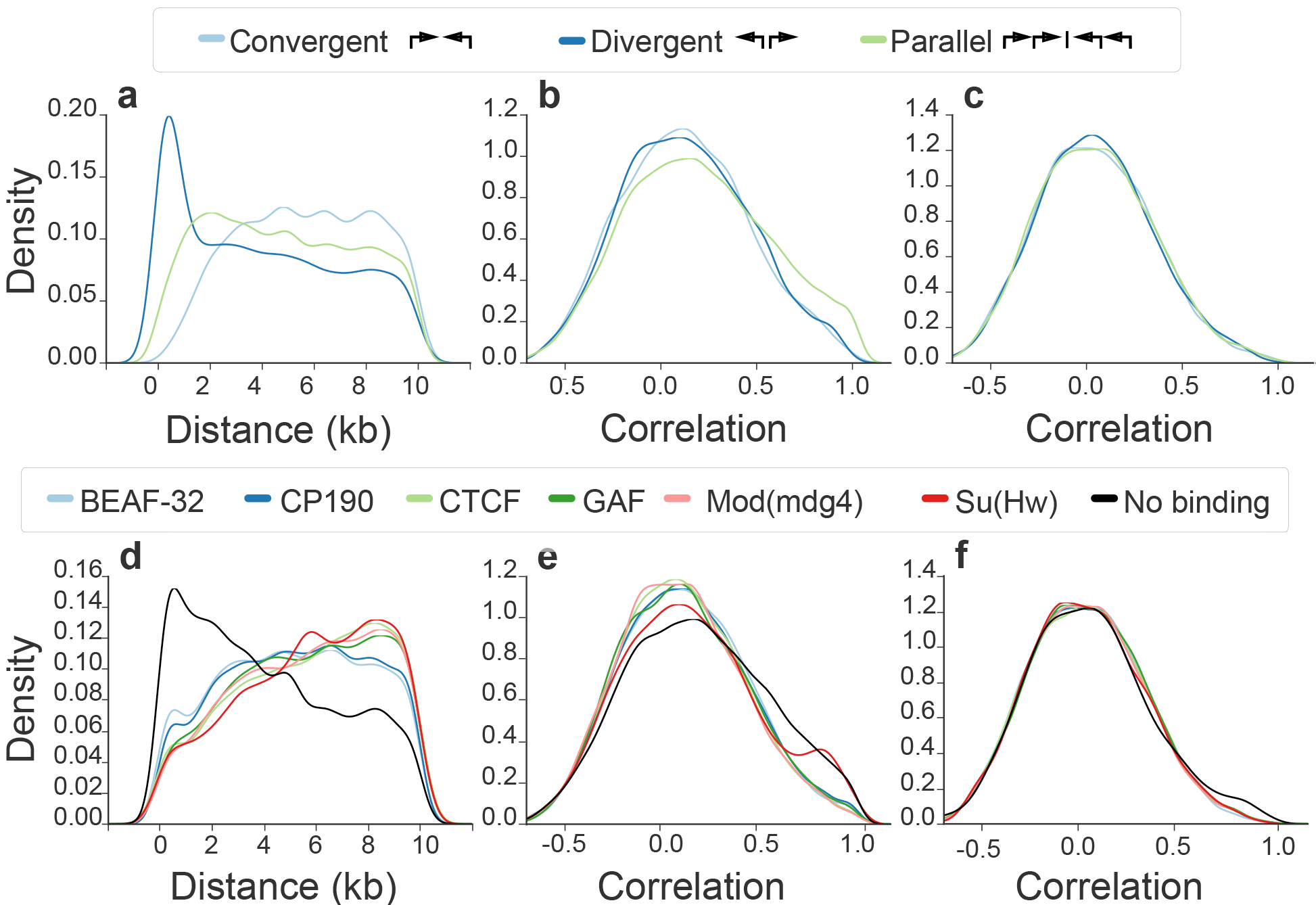
EP distance causes gene orientation-dependent correlation and provides regulatory independence to gene neighbors. Distribution of intergenic distances below 10 kb of gene pairs in *D. melanogaster* by configuration (~5 to 18 thousand gene pairs for each group, a) and flanking insulator binding sites identified through ChIP-chip (Negre et al., 2010) (~5 to 15 thousand pairs for each group, d). The corresponding distribution of correlations is shown for the same gene pairs (b, e) and pairs with controlled distributions of intergenic distances between 30 and 40 kb (~7 to 14 thousand pairs for gene orientation groups, ~10 to 18 thousand for insulator groups, c, f).

From the regulatory perspective, EP distance provides independence to most gene pairs, as the vast majority have an intergenic distance that puts them in the baseline correlation regime (Figure 1c). To study the enhancer-blocking influence of insulators (Bushey et al., 2009) genome-wide, we analyzed each group of genes flanked by insulator binding sites, which were previously determined using Chromatin ImmunoPrecipitation coupled with microarrays (ChIP-chip) for the six known insulators in *D. melanogaster*: BEAF-32, CP190, CTCF, GAF, Mod(mdg4) and Su(Hw) (Negre et al., 2010). We observed that gene neighbors closer than 10 kb bound by each of the insulators tend to be less correlated in gene expression than gene pairs not bound by them (Figure 3e), supporting their role in genome-wide insulation and agreeing with the observation that insulators tend to separate differentially expressed genes (Negre et al., 2010; Xie et al., 2007). Nevertheless, the same groups of gene pairs also tend to have much larger intergenic distances than genes that are not flanked by insulator binding sites (Figure 3d). After controlling for the distribution of intergenic distances, we found very similar correlation distributions between insulator and not insulator flanked gene pairs (Figure 3f). This agrees with previous reports suggesting that insulators do not block enhancers everywhere they bind, but rather act only on very specific genomic contexts (Liu et al., 2015; Schwartz et al., 2012); it also reconciles the lack of known insulator orthologs in *C. elegans* (Heger et al., 2009) in the context of local enhancer-blocking. In combination, these studies strongly suggest that EP distance is the general source of transcriptional independence for close gene neighbors.

Previous EP compatibility studies suggest that EP specificity is widespread (Gehrig et al., 2009), while others that it is restricted to a smaller subset of enhancers (Butler and Kadonaga, 2001). Although our results support the latter, views arising from these studies might not be mutually exclusive, as it is likely that enhancers have specificity to promoter classes (Danino et al., 2015), whose limited number could result in general EP compatibility. Nevertheless, the implications from considering our observations are broadly applicable to gene regulation and likely act in conjunction with other regulatory mechanisms, such as chromatin accessibility. For example, position effects, in which transgene expression levels are influenced by the insertion site (Gierman et al., 2007), are naturally expected from enhancer sharing. Chromosomal translocations and mutations involving regulatory elements likely impact genetic contexts rather than individual genes. Furthermore, enhancer scaling by EP distance contributes to the weight of any given EP interaction and thus enriches both the complexity and tunability of genomic logic. Our analysis provides a clarifying perspective of gene regulation consistent with both mechanistic and genome-wide studies.

## Experimental procedures

### Computational biology

For each analyzed organism, Ensembl (Flicek et al., 2014) protein-coding genes were grouped by chromosome, sorted by position, and paired with the 100 nearest neighbors within the same chromosome. A list of duplicated gene pairs for *H. sapiens* and *M. musculus* was obtained from the Duplicated Genes Database (Ouedraogo et al., 2012) (http://dgd.genouest.org). A list of *C. elegans* genes predicted to be in operons was obtained from Allen et al. (2011), and gene pairs present in the same operon were removed from the analysis to prevent co-transcriptional bias. Processed RNA-seq data was obtained from multiple sources (Attrill et al., 2016; Ellahi et al., 2015; Gerstein et al., 2010; The ENCODE Project Consortium, 2012) and converted to transcripts per million (TPM) (Wagner et al., 2012) when necessary. Formatted datasets are available upon request. Genes detected in less than 80% of experiments were discarded. To compute the Spearman correlation coefficient, TPM values were ranked in each RNA-seq experiment and the pairwise Pearson correlation coefficient was computed on the ranked values according to the following equation:

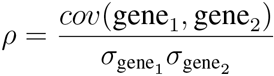

where gene_1_ and gene_2_ are the corresponding ranks of each paired gene in a given RNA-seq experiment, *cov* their covariance and *σ* their standard deviation. The list of gene pairs with intergenic distances and correlation coefficients was sorted by increasing intergenic distance, and subsequently smoothed using a sliding median with window size of 1000 gene pairs. The result was then fitted to an exponential decay function of the form:

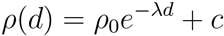

where *ρ*_0_ is the median Spearman correlation coefficient of the closest neighboring genes, *d* the intergenic distance and *c* the baseline correlation. The mean distance at which a pair of genes remain correlated was then computed as:

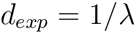

A list of genes annotated with RNA *in situ* hybridization data (Hammonds et al., 2013; Tomancak et al., 2002, 2007) was obtained from the Berkeley Drosophila Genome Project (http://insitu.fruit-fly.org). Insulator ChIP-chip data was obtained from Negre et al. (2010) (GSE16245); the intersection of replicates was used. HiC data was obtained from Rao et al. (2014) (GSE63525, GM12878 primary replicate HiCCUPS looplist). Functional protein classification was conducted using Panther (Mi et al., 2016). Genomic manipulations were conducted using Bedtools v2.24.0 (Quinlan and Hall, 2010). Data analysis was conducted using Python 2.7.9 and the Scipy library (McKinney, 2010).

### Molecular biology

*C. elegans* was cultured under standard laboratory conditions (Stiernagle, 2006). For enhancer additivity experiments, transgenic *C. elegans* lines carrying extra-chromosomal arrays were generated by injecting each plasmid at 50 ng/μl into *unc-119* mutant animals. The minimal Δ*pes-10* promoter (Fire et al., 1990) was used in all constructs. Minimal regions of the *myo-2* and *unc-54* enhancers (Okkema et al., 1993) able to drive tissue specific expression were used. The BWM enhancer was obtained from the upstream region of *F44B9.2*; the BWM plus intestine enhancer was obtained from the upstream region of *rps-1*. Animals were imaged at 40x using a GFP filter on a Zeiss Axioskop microscope. For the enhancer promoter distance and ectopic enhancer experiments, we defined an EP distance of 0 to be the enhancer placed just upstream of the Δ*pes-10* promoter, which is ~350 bp. To ensure neutrality yet maintain a similar GC content as non-coding sequences in *C. elegans*, we used non-overlapping AT-rich DNA spacers obtained from the genome of *Escherichia coli*. Constructs were integrated in single-copy into chromosome IV via CRISPR-Cas9 using plasmids provided as gifts by Dr. Zhiping Wang and Dr. Yishi Jin (unpublished results). Briefly, plasmids containing the following expresssion cassettes were co-injected: reporter and hygromycin resistance genes flanked by homologous arms for recombination-directed repair (10ng/μl), single-guide RNA (10ng/μl), Cas9 (10ng/μl), and extra-chromosomal array reporter for expression of either *rfp* or *gfp* outside the tissue of interest (10ng/μl). Transformants were selected for using hygromycin at 10 μg/μl, and those not carrying extra-chromosomal transgenes, lacking of *gfp* or *rfp* expression outside the tissue of interest, were subsequently isolated. Animals homozygotic for the insertion were identified by polymerase-chain reaction (PCR) and Sanger sequencing. Quantitative PCR was carried out as previously described (Ly et al., 2015) using *pmp-3* as a reference gene (Zhang et al., 2012). Briefly, third-stage larval (L3) worms, when expression from the test enhancers is maximal according to RNA-seq data, were synchronized at 20°C via egg-laying. 15 animals were lysed in 1.5μl of Lysis Buffer (5 mM Tris pH 8.0 (MP Biomedicals), 0.5% Triton X-100, 0.5% Tween 20, 0.25 mM EDTA (Sigma-Aldrich)) with proteinase-K (Roche) at 1.5 μg/μl, incubated at 65°C for 10 minutes followed by 85°C for one minute. Reverse transcription was carried out using the Maxima H Minus cDNA synthesis kit (Thermo Fisher) by adding 0.3 μl H_2_O, 0.6 μl 5x enzyme buffer, 0.15 μl 10mM dNTP mix, 0.15 μl 0.5 μg/μl oligo dT primer, 0.15μl enzyme mix and 0.15 μl DNAse, and incubated for 2 minutes at 37°C, followed by 30 minutes at 50°C and finally 2 minutes at 85°C. The cDNA solution was diluted to 15 μl and 1 μl was used for each qPCR reaction, so that on average each well contained the amount of RNA from a single worm. All qPCR reactions were performed with three technical replicates and at least three biological replicates using the Roche LightCycler^®^ 480 SYBR Green I Master in the LightCycler^®^ 480 System. Crossing point-PCR-cycle (Cp) averages were computed for each group of three technical replicates; these values were then substracted from the respective average Cp value of the reference gene.

### Data and reagent availability

Strains are available upon request. Relevant DNA sequences, including spacers, enhancers, primers, sgRNA and homology arms are available in the supplementary table 1. Correlation datasets are available in the supplementary file 1. Python source code, and links to all expression datasets used in this study, are available for download on the following github repository: https://github.com/WormLabCaltech/QuinteroSternberg2016.git.

## Acknowledgments

Our work was supported by the Howard Hughes Medical Institute, of which P.W.S is an investigator. We thank Zhiping Wang and Yishi Jin for plasmids for Crispr-Cas9 single copy insertion, Carmie Robinson for discussions, experimental suggestions and comments on the manuscript, Han Wang for discussions, technical advise and comments on the manuscript, Hillel Schwartz, Mitchell Guttman, Mihoko Kato, David Angeles-Albores, Jonathan Liu, Barbara Wold, Isabelle Peter and Angelike Stathopoulos for discussions and comments on the manuscript, the Encode and ModEncode consortiums, Flybase, Wormbase and Ensembl databases, the Wold Lab and the Guigo Lab for data accessibility. This paper is dedicated to the memory of Eric H. Davidson.

## Contributions

P.Q.C. performed the experiments and analyzed the data. P.Q.C. and P.W.S. designed the experiments and wrote the paper.

---

**Supplementary Figure 1:**
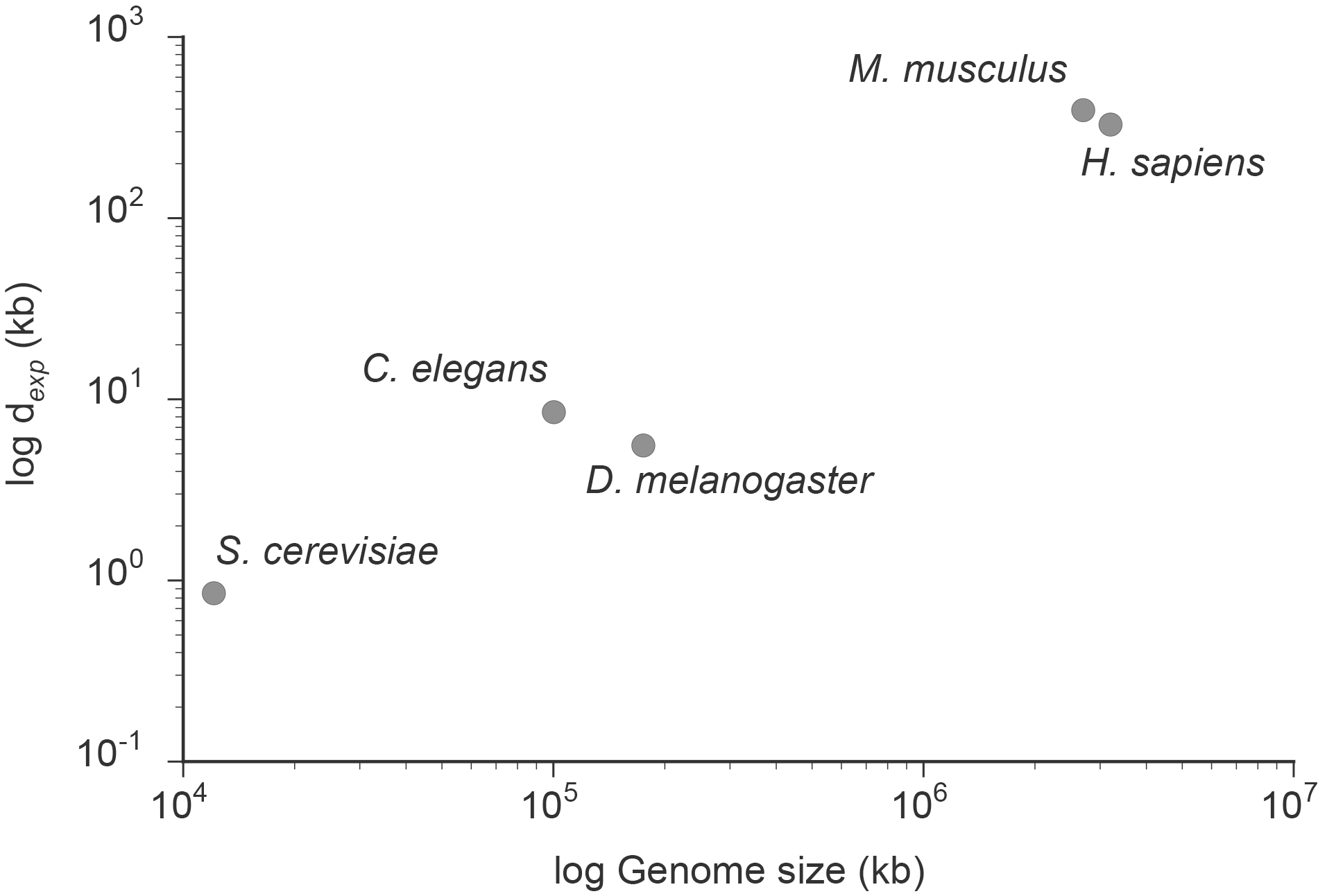
The distance at which a pair of genes remain correlated (*d_exp_*) scales with genome size.

**Supplementary Figure 2:**
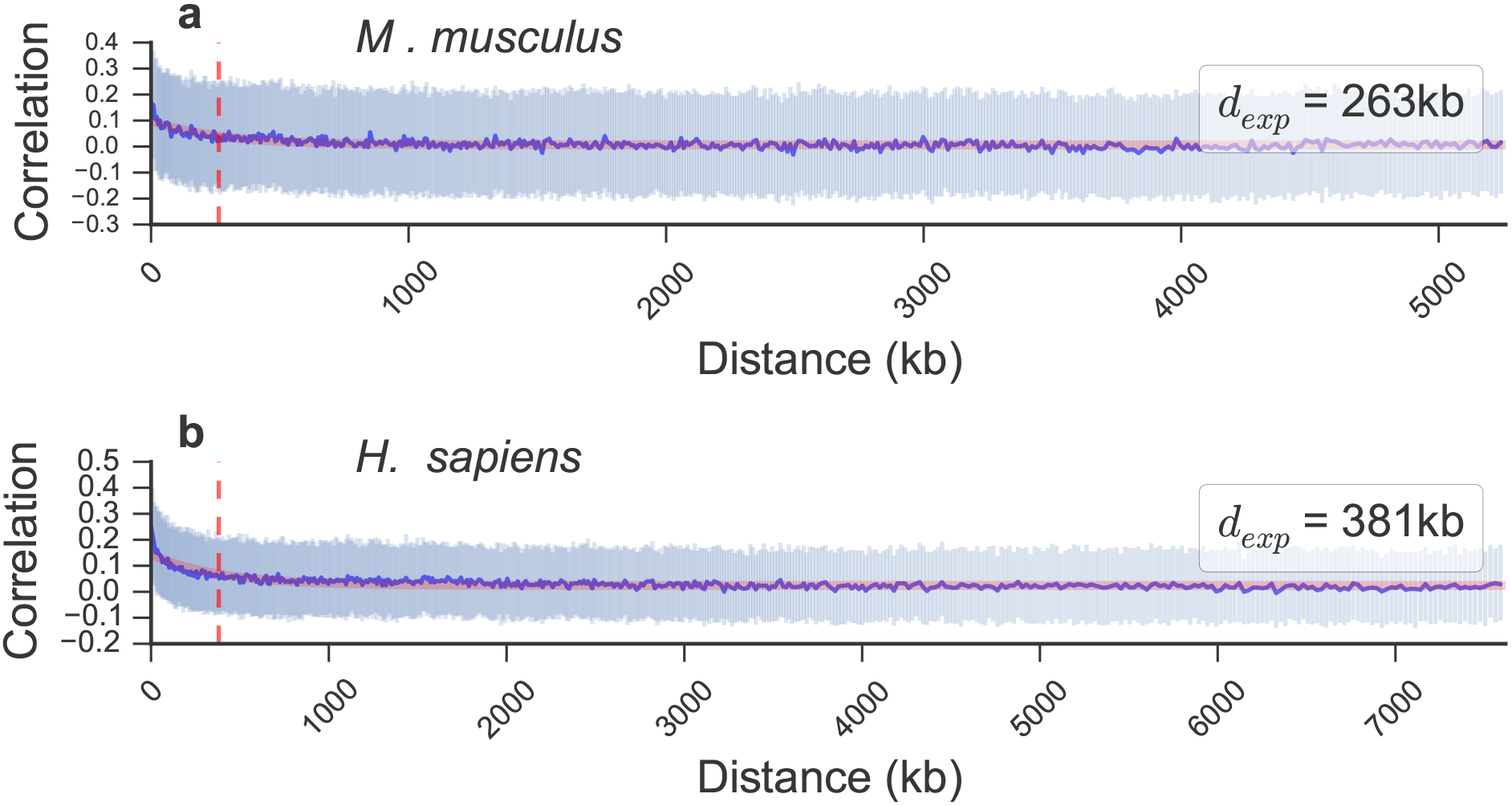
Removing duplicated genes does not affect the overall correlation of gene neighbors. Sliding median of correlations between paired neighbors (blue line) and interquartile range (pale blue) with increasing intergenic distance in *M. musculus* (a) and *H. sapiens* (b) after removing duplicated gene pairs. Fit to exponential decay function (red line) and corresponding *d_exp_* (red dashed line) are shown.

**Supplementary Figure 3:**
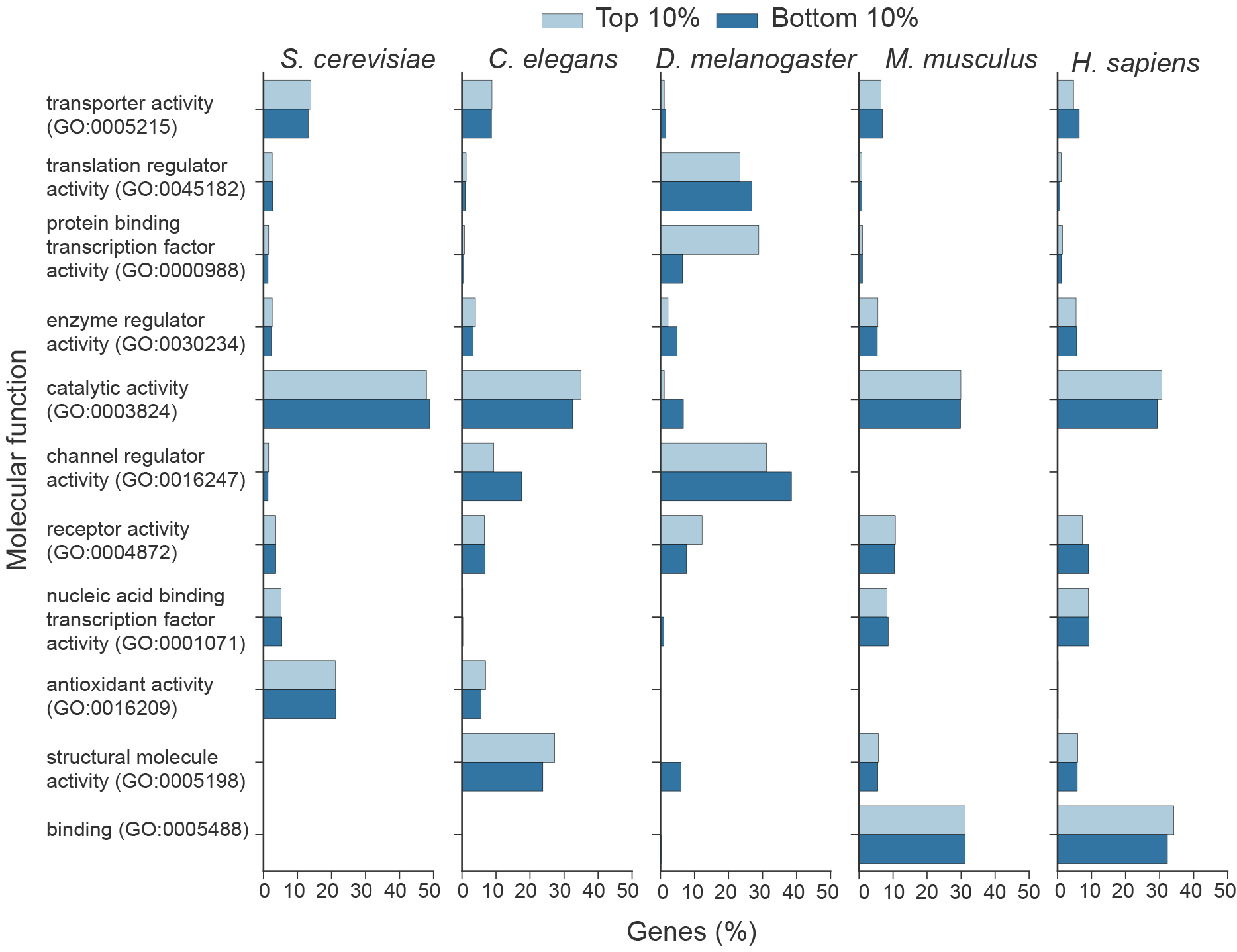
Representation of gene ontology annotations remains unbiased in correlated gene pairs. The molecular function classification of top and bottom 10% correlated gene pairs with intergenic distance below *d_exp_* is shown for each organism.

**Supplementary Figure 4:**
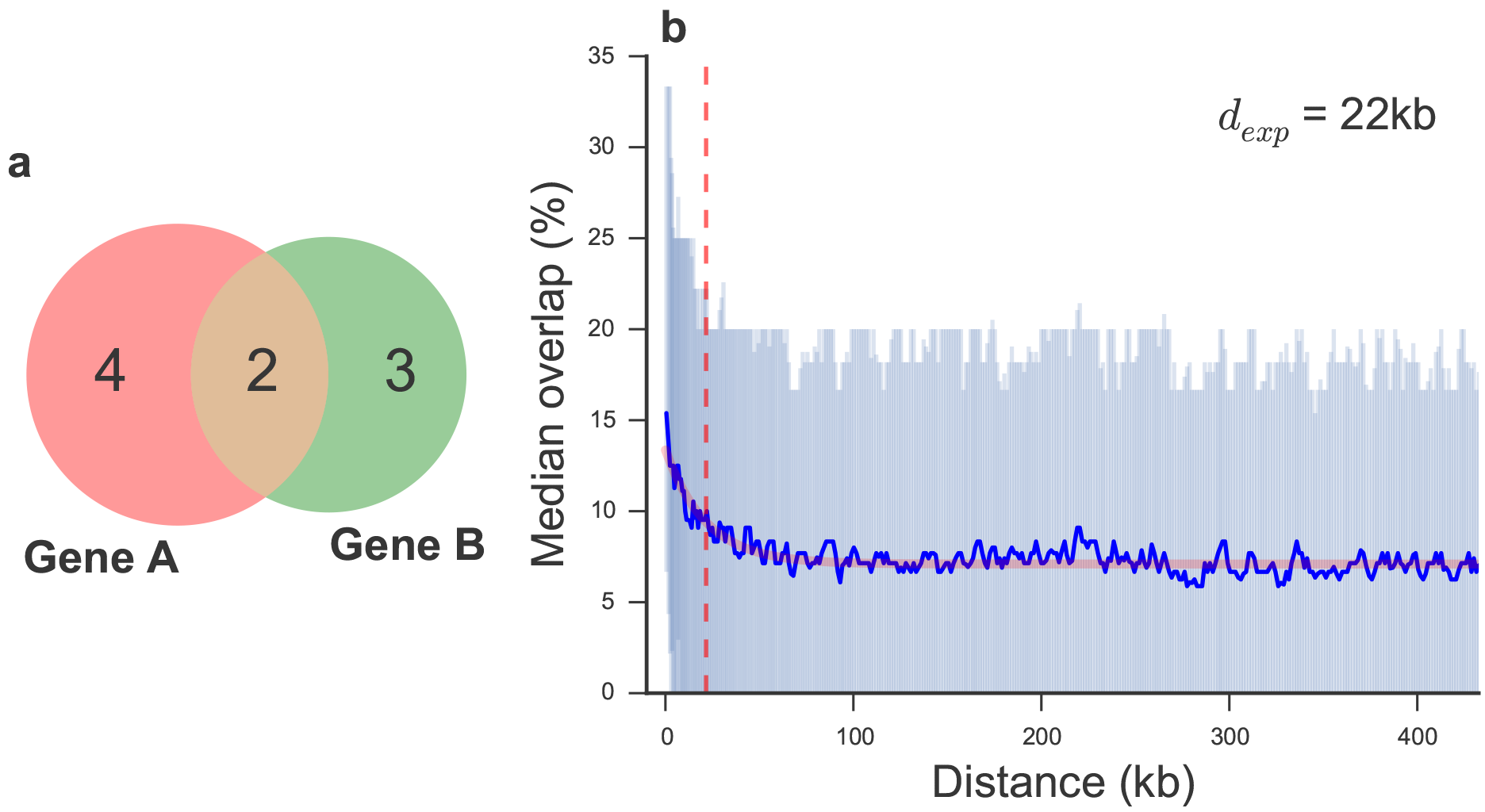
Gene pairs are correlated in spatial expression in *D. melanogaster*. The size of the intersection between the set of tissues in which each gene of a given pair is expressed was divided over the size of the union of the same sets. An example is shown in (a), where the percentage overlap is 2/(4+3−2)=0.4. b) Sliding median of the percentage overlap in tissue specific expression (blue line) and interquartile range (pale blue) with increasing intergenic distance. Fit to exponential decay function (red line) and corresponding *d_exp_* (red dashed line) are shown.

**Supplementary Figure 5:**
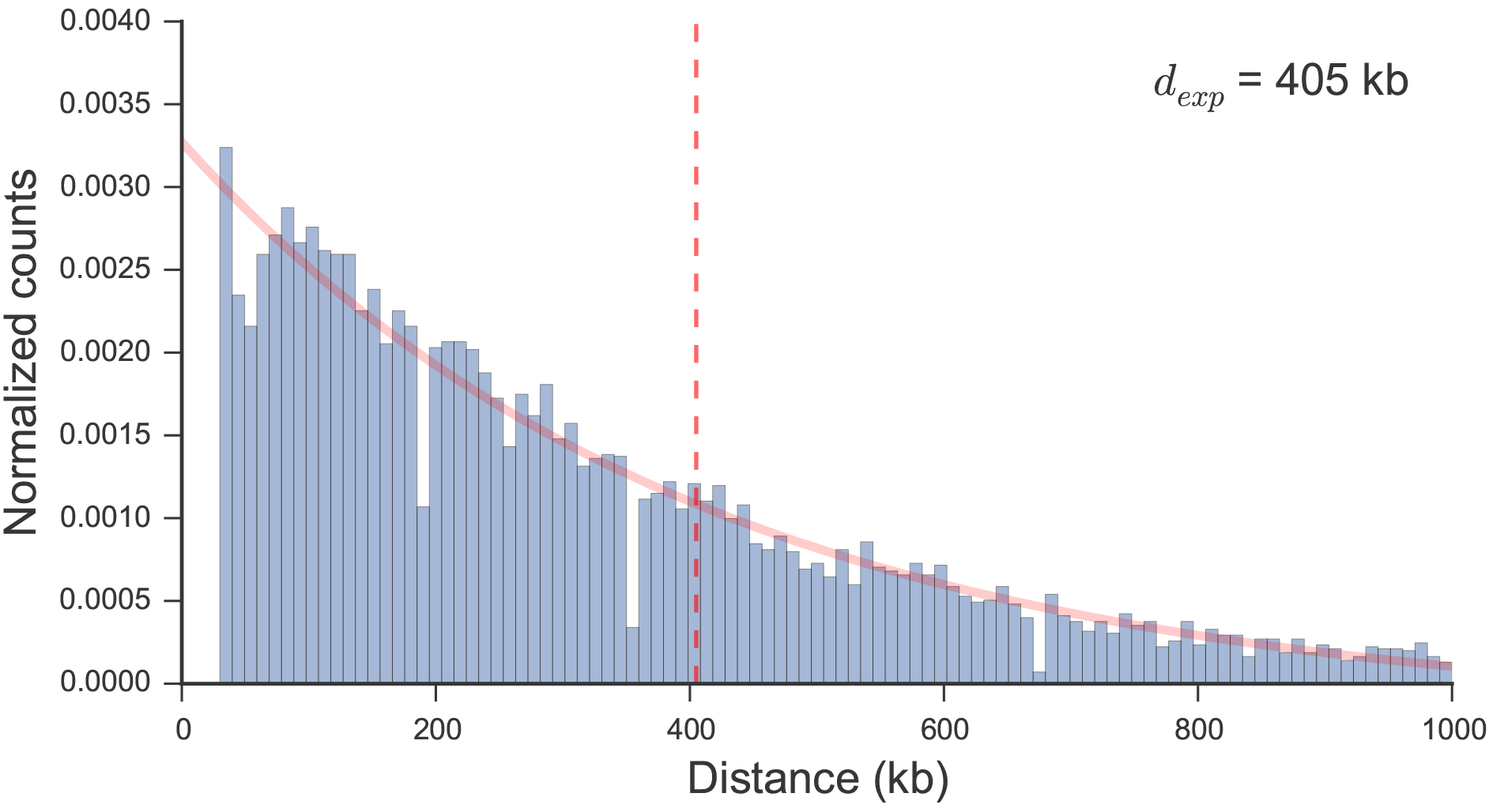
Chromatin looping decreases exponentially with distance in human cell lines. Normalized count of loops identified through HiC by Rao et al. (2014) were fit to exponential decay function (red line); the resulting *d_exp_* (red dashed line) is shown.

**Supplementary Figure 6:**
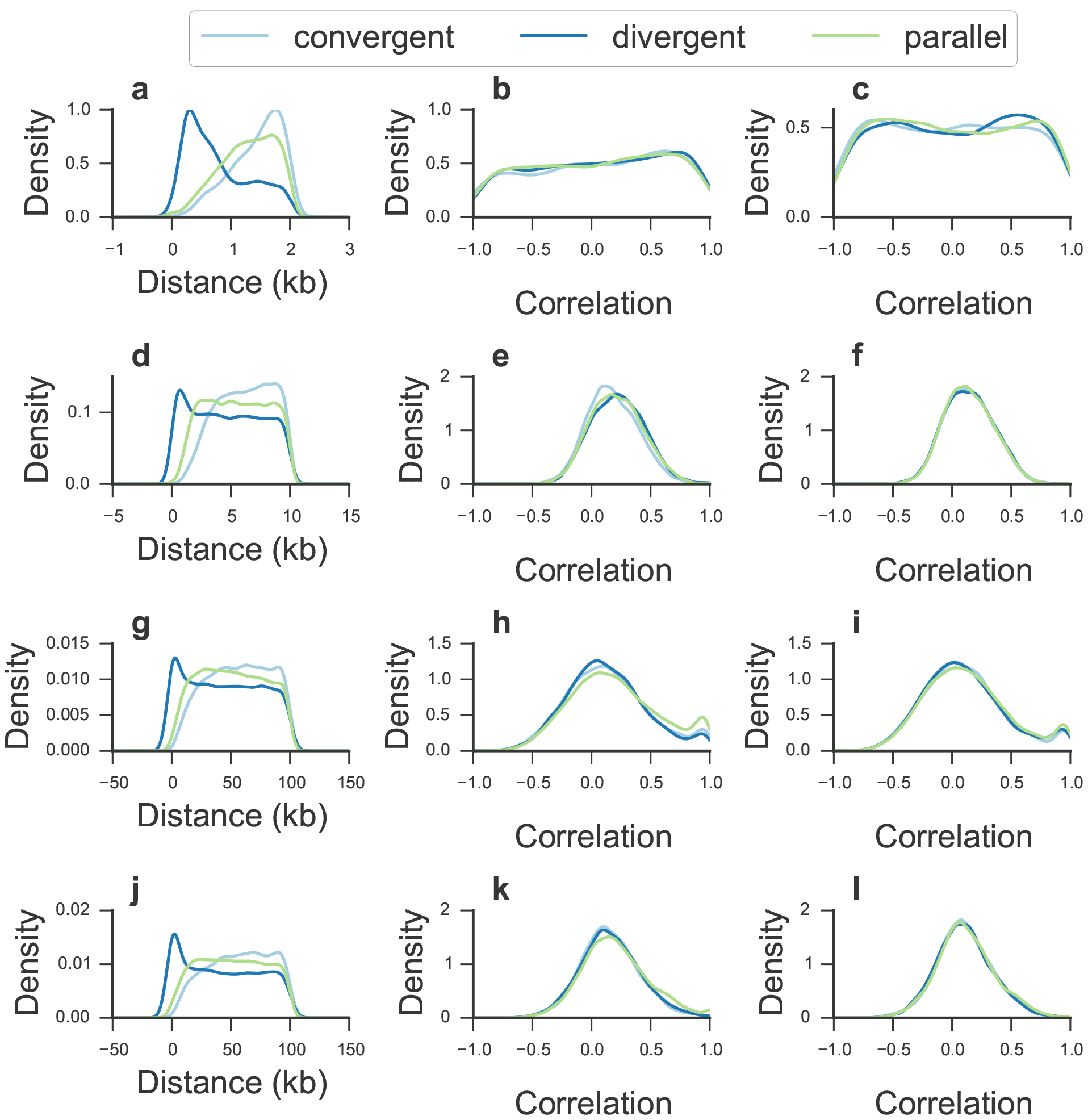
Gene orientation effect in correlation of gene pairs is explained by EP distance. Distribution of intergenic distances and the corresponding distribution of correlations of gene pairs is shown in the first and second columns, respectively; correlations after controlling for intergenic distance are shown in the third column. The range of distances between paired genes for each plot is as follows: *S. cerevisiae* below 2 kb (a,b) and between 2 and 4 kb (c). *C. elegans* below 10 kb (d,e) and between 10 and 20 kb (f). *H. sapiens* and *M. musculus* below 100 kb (g, h, j, k) and between 100 and 200 kb (i, l).

## References

Allen, M. A., Hillier, L. W., Waterston, R. H., and Blumenthal, T. (2011). A global analysis of C. elegans trans-splicing. Genome Res., 21:255–264.

Araya, C. L., Kawli, T., Kundaje, A., Jiang, L., Wu, B., Vafeados, D., Terrell, R., Weissdepp, P., Gevirtzman, L., Mace, D., Niu, W., Boyle, A. P., Xie, D., Ma, L., Murray, J. I., Reinke, V., Waterston, R. H., and Snyder, M. (2014). Regulatory analysis of the C. elegans genome with spatiotemporal resolution. Nature, 512(7515):400-405.

Attrill, H., Falls, K., Goodman, J. L., Millburn, G. H., Antonazzo, G., Rey, A. J., Marygold, S. J., and the FlyBase consortium (2016). Flybase: establishing a gene group resource for drosophila melanogaster. Nucleic Acids Res., 44(D1):D786-D792.

Bushey, A. M., Dorman, E. R., and Corces, V. G. (2009). Chromatin insulators:regulatory mechanisms and epigenetic inheritance. Mol. Cell, 32(404):1-9.

Butler, J. E. and Kadonaga, J. T. (2001). Enhancer-promoter specificity mediated by dpe or tata core promoter motifs. Genes Development, 15(19):2515–2519.

Chen, N. and Stein, L. D. (2006). Conservation and functional significance of gene topology in the genome of Caenorhabditis elegans. Genome Res., 16(5):606–617.

Corsi, A. K., Wightman, B., and Chalfie, M. (2015). A transparent window into biology: A primer on Caenorhabditis elegans. Genetics, 200(2):387–407.

Danino, Y. M., Even, D., Ideses, D., and Juven-Gershon, T. (2015). The core promoter: At the heart of gene expression. Biochimica et Biophysica Acta (BBA) - Gene Regulatory Mechanisms, 1849(8):1116–1131.

Davidson, E. H. and Peter, I. S. (2015). Chapter 1 - the genome in development. In Davidson, E. H. and Peter, I. S., editors, Genomic Control Process, pages 1-40. Academic Press, Oxford.

Dupuy, D., Li, Q.-R., Deplancke, B., Boxem, M., Hao, T., Lamesch, P., Sequerra, R., Bosak, S., Doucette-Stamm, L., Hope, I. A., Hill, D. E., Walhout, A. J., and Vidal, M. (2004). A first version of the caenorhabditis elegans promoterome. Genome Res., 14(10b):2169-2175.

Ellahi, A., Thurtle, D. M., and Rine, J. (2015). The chromatin and transcriptional landscape of native saccharomyces cerevisiae telomeres and subtelomeric domains. Genetics, 2:505–521.

Fire, A., Harrison, S. W., and Dixon, D. (1990). A modular set of lacz fusion vectors for studying gene expression in caenorhabditis elegans. Gene, 93(2):189–198.

Flicek, P., Amode, M. R., Barrell, D., Beal, K., Billis, K., Brent, S., Carvalho-Silva, D., Clapham, P., Coates, G., Fitzgerald, S., Gil, L., Giron, C. G., Gordon, L., Hourlier, T., Hunt, S., Johnson, N., Juettemann, T., Kahari, A. K., Keenan, S., Kulesha, E., Martin, F. J., Maurel, T., McLaren, W. M., Murphy, D. N., Nag, R., Overduin, B., Pignatelli, M., Pritchard, B., Pritchard, E., Riat, H. S., Ruffier, M., Sheppard, D., Taylor, K., Thormann, A., Trevanion, S. J., Vullo, A., Wilder, S. P., Wilson, M., Zadissa, A., Aken, B. L., Birney, E., Cunningham, F., Harrow, J., Herrero, J., Hubbard, T. J., Kinsella, R., Muffato, M., Parker, A., Spudich, G., Yates, A., Zerbino, D. R., and Searle, S. M. (2014). Ensembl 2014. Nucleic Acids Res., 42(D1):D749-D755.

Gehrig, J., Reischl, M., Kalmar, E., Ferg, M., Hadzhiev, Y., Zaucker, A., Song, C., Schindler, S., Liebel, U., and Muller, F. (2009). Automated high-throughput mapping of promoter-enhancer interactions in zebrafish embryos. Nat Meth, 6(12):911–916.

Gerstein, M. B., Lu, Z. J., Van Nostrand, E. L., Cheng, C., Arshinoff, B. I., Liu, T., Yip, K. Y., Robilotto, R., Rechtsteiner, A., Ikegami, K., Alves, P., Chateigner, A., Perry, M., Morris, M., Auerbach, R. K., Feng, X., Leng, J., Vielle, A., Niu, W., Rhrissorrakrai, K., Agarwal, A., Alexander, R. P., Barber, G., Brdlik, C. M., Brennan, J., Brouillet, J. J., Carr, A., Cheung, M.-S., Clawson, H., Contrino, S., Dannenberg, L. O., Dernburg, A. F., Desai, A., Dick, L., Dose, A. C., Du, J., Egelhofer, T., Ercan, S., Euskirchen, G., Ewing, B., Feingold, E. A., Gassmann, R., Good, P. J., Green, P., Gullier, F., Gutwein, M., Guyer, M. S., Habegger, L., Han, T., Henikoff, J. G., Henz, S. R., Hinrichs, A., Holster, H., Hyman, T., Iniguez, A. L., Janette, J., Jensen, M., Kato, M., Kent, W. J., Kephart, E., Khivansara, V., Khurana, E., Kim, J. K., Kolasinska-Zwierz, P., Lai, E. C., Latorre, I., Leahey, A., Lewis, S., Lloyd, P., Lochovsky, L., Lowdon, R. F., Lubling, Y., Lyne, R., MacCoss, M., Mackowiak, S. D., Mangone, M., McKay, S., Mecenas, D., Merrihew, G., Miller, D. M., Muroyama, A., Murray, J. I., Ooi, S.-L., Pham, H., Phippen, T., Preston, E. A., Rajewsky, N., Ratsch, G., Rosenbaum, H., Rozowsky, J., Rutherford, K., Ruzanov, P., Sarov, M., Sasidharan, R., Sboner, A., Scheid, P., Segal, E., Shin, H., Shou, C., Slack, F. J., Slightam, C., Smith, R., Spencer, W. C., Stinson, E. O., Taing, S., Takasaki, T., Vafeados, D., Voronina, K., Wang, G., Washington, N. L., Whittle, C. M., Wu, B., Yan, K.-K., Zeller, G., Zha, Z., Zhong, M., Zhou, X., Ahringer, J., Strome, S., Gunsalus, K. C., Micklem, G., Liu, X. S., Reinke, V., Kim, S. K., Hillier, L. W., Henikoff, S., Piano, F., Snyder, M., Stein, L., Lieb, J. D., and Waterston, R. H. (2010). Integrative analysis of the Caenorhabditis elegans genome by the modencode project. Science, 330(6012):1775–1787.

Ghanbarian, A. T. and Hurst, L. D. (2015). Neighboring genes show correlated evolution in gene expression. Mol. Biol. Evol., 32(7):1748–1766.

Gierman, H. J., Indemans, M. H., Koster, J., Goetze, S., Seppen, J., Geerts, D., van Driel, R., and Versteeg, R. (2007). Domain-wide regulation of gene expression in the human genome. Genome Res., 17(9):1286–1295.

Hammonds, A. S., Bristow, C. A., Fisher, W. W., Weiszmann, R., Wu, S., Hartenstein, V., Kellis, M., Yu, B., Frise, E., and Celniker, S. E. (2013). Spatial expression of transcription factors in drosophila embryonic organ development. Genome Biol., 14(12):R140.

He, B., Chen, C., Teng, L., and Tan, K. (2014). Global view of enhancer-promoter interactome in human cells. Proc. Natl. Acad. Sci. USA, 111(21):E2191-E2199.

Heger, P., Marin, B., and Schierenberg, E. (2009). Loss of the insulator protein CTCF during nematode evolution. BMC Mol. Biol., 5:1–14.

Kagey, M. H., Newman, J. J., Bilodeau, S., Zhan, Y., Orlando, D. A., van Berkum, N. L., Ebmeier, C. C., Goossens, J., Rahl, P. B., Levine, S. S., Taatjes, D. J., Dekker, J., and Young, R. A. (2010). Mediator and cohesin connect gene expression and chromatin architecture. Nature, 467(7314):430–435.

Liu, M., Maurano, M. T., Wang, H., Qi, H., Song, C.-z., Navas, P. A., Emery, D. W., Stamatoyannopoulos, J. A., and Stamatoyannopoulos, G. (2015). Genomic discovery of potent chromatin insulators for human gene therapy. Nat. biotechnol., 33(2):198–203.

Ly, K., Reid, S. J., and Snell, R. G. (2015). Rapid RNA analysis of individual Caenorhabditis elegans. MethodsX, 2:59–63.

Malik, S. and Roeder, R. G. (2010). The metazoan mediator co-activator complex as an integrative hub for transcriptional regulation. Nat. Rev. Genet., 11(11):761–772.

Marsman, J. and Horsfield, J. A. (2012). Long distance relationships: Enhancer-promoter communication and dynamic gene transcription. Biochimica et Biophysica Acta (BBA) - Gene Regulatory Mechanisms, 1819(11-12):1217-1227.

McKinney, W. (2010). Data structures for statistical computing in python. In van der Walt, S. and Millman, J., editors, Proceedings of the 9th Python in Science Conference, pages 51-56.

Mi, H., Poudel, S., Muruganujan, A., Casagrande, J. T., and Thomas, P. D. (2016). Panther version 10: expanded protein families and functions, and analysis tools. Nucleic Acids Res., 44(D1):D336-D342.

Michalak, P. (2008). Coexpression, coregulation, and cofunctionality of neighboring genes in eukaryotic genomes. Genomics, 91(3):243–248.

Negre, N., Brown, C. D., Shah, P. K., Kheradpour, P., Morrison, C. A., Henikoff, S., Kellis, M., and White, K. P. (2010). A Comprehensive Map of Insulator Elements for the Drosophila Genome. Plos Genet., 6(1):e1000814.

Okkema, P. G., Harrison, S. W., Plunger, V., Aryana, A., and Fire, A. (1993). Sequence Requirements for Myosin Gene Expression and Regulation in C elegans. Genetics, 135(Waterston 1988):385–404.

Ouedraogo, M., Bettembourg, C., Bretaudeau, A., Sallou, O., Diot, C., Demeure, O., and Lecerf, F. (2012). The duplicated genes database: Identification and functional annotation of co-localised duplicated genes across genomes. PLoS ONE, 7(11):1–8.

Quinlan, A. R. and Hall, I. M. (2010). Bedtools: a flexible suite of utilities for comparing genomic features. Bioinformatics, 26(6):841–842.

Rao, S. S. P., Huntley, M. H., Durand, N. C., Stamenova, E. K., Bochkov, I. D., Robinson, J. T., Sanborn, A. L., Machol, I., Omer, A. D., Lander, E. S., and Aiden, E. L. (2014). A 3d map of the human genome at kilobase resolution reveals principles of chromatin looping. Cell, 159(7):1665–1680.

Ringrose, L., Chabanis, S., Angrand, P. O., Woodroofe, C., and Stewart, A. F. (1999). Quantitative comparison of dna looping in vitro and in vivo: chromatin increases effective dna flexibility at short distances. EMBO J., 18(23):6630–6641.

Schwartz, Y. B., Linder-basso, D., Kharchenko, P. V., Tolstorukov, M. Y., Kim, M., Li, H.-b., Gorchakov, A. A., Minoda, A., Shanower, G., Alekseyenko, A. A., Riddle, N. C., Jung, Y. L., Gu, T., Plachetka, A., Elgin, S. C. R., Kuroda, M. I., Park, P. J., Savitsky, M., and Karpen, G. H. (2012). Nature and function of insulator protein binding sites in the Drosophila genome. Genome Res., 11:2188–2198.

Stiernagle, T. (2006). Maintenance of c. elegans. In C. elegans Research Community, editor, WormBook. WormBook. The ENCODE Project Consortium (2012). An integrated encyclopedia of dna elements in the human genome. Nature, 489(7414):57–74.

Tomancak, P., Beaton, A., Weiszmann, R., Kwan, E., Shu, S., Lewis, S. E., Richards, S., Ash-burner, M., Hartenstein, V., Celniker, S. E., and Rubin, G. M. (2002). Systematic determination of patterns of gene expression during drosophila embryogenesis. Genome Biol., 3(12):research0088.1–88.14.

Tomancak, P., Berman, B. P., Beaton, A., Weiszmann, R., Kwan, E., Hartenstein, V., Celniker, S. E., and Rubin, G. M. (2007). Global analysis of patterns of gene expression during drosophila embryogenesis. Genome Biol., 8(7):R145.

van Arensbergen, J., van Steensel, B., and Bussemaker, H. J. (2014). In search of the determinants of enhancer-promoter interaction specificity. Trends in cell biology, 24(11):695–702.

Wagner, G. P., Kin, K., and Lynch, V. J. (2012). Measurement of mrna abundance using rna-seq data: Rpkm measure is inconsistent among samples. Theory Biosci., 131(4):281–285.

Xie, X., Mikkelsen, T. S., Gnirke, A., Lindblad-toh, K., Kellis, M., and Lander, E. S. (2007). Systematic discovery of regulatory motifs in conserved regions of the human genome, including thousands of CTCF insulator sites. Proc. Natl. Acad. Sci. USA, 104(17):7145–7150.

Zhang, Y., Chen, D., Smith, M. A., Zhang, B., and Pan, X. (2012). Selection of reliable reference genes in /textitCaenorhabditis elegans for analysis of nanotoxicity. PLoS ONE, 7(3):1–7.

